# Survival and ice nucleation activity of *Pseudomonas syringae* strains exposed to simulated high-altitude atmospheric conditions

**DOI:** 10.1101/408906

**Authors:** Gabriel Guarany de Araujo, Fabio Rodrigues, Fabio Luiz Teixeira Gonçalves, Douglas Galante

## Abstract

The epiphytic bacterium *Pseudomonas syringae* produces the most efficient and well-studied biological ice nuclei (IN) known. Bioaerosols containing these cells have been proposed to influence cloud glaciation, an important process in the initiation of precipitation. The presence of this species has been reported on rain, snow, and cloud water samples, but how these organisms can survive the harsh conditions present on the high atmosphere still remains to be better understood. In this study, the impact caused by this type of environment on *P. syringae* was assayed by measuring their viability and IN activity. Two strains, of the pathovars *syringae* and *garcae*, were compared to *Escherichia coli.* While UV-C radiation effectively inactivated these cells, the *Pseudomonas* were much more tolerant to UV-B. The *P. syringae* strains were also more resistant to “environmental” UV radiation from a solar simulator, composed of UV-A and UV-B. The response of their IN after long exposures to this radiation varied: only one strain suffered a relatively small 10-fold reduction in IN activity at -5 °C. Desiccation at different relative humidity values also affected the IN, but some activity at -5 °C was still maintained for all tests. The pathovar *garcae* tended to be more resistant to the stress treatments than the pathovar *syringae*, particularly to desiccation, though its IN were found to be more sensitive. Compared to *E. coli*, the *P. syringae* strains seemed relatively better adapted to survival under conditions present on the atmosphere at high altitudes.

**IMPORTANCE:** The plant-associated bacterium *Pseudomonas syringae* produces on its outer membrane highly efficient ice nuclei which are able to induce the freezing of supercooled water. This ability has been linked to increased frost damaged on colonized leaves and also to the formation of ice in clouds, an important process leading to precipitation. *P. syringae* has been found on rain, snow, and cloud water samples, confirming its presence on the atmosphere. This study aimed to assess the survival of these cells and the maintenance of their ice nucleation activity under stressing conditions present in high altitudes: ultraviolet radiation and desiccation. *P. syringae* strains were shown to at least partially tolerate these factors, and their most efficient ice nuclei, while affected, could still be detected after all experiments.

## INTRODUCTION

The Gram-negative bacterium *Pseudomonas syringae* is a common member of epiphytic communities and an important phytopathogen in diverse crops (Hirano and Upper, 2000). It was the first organism found to produce biological ice nuclei (IN) of exceptional efficiency, being able to freeze supercooled water at temperatures above -10 °C (Maki *et al.*, 1974). The IN activity of this organism originates from a large protein (called InaZ) situated at the cell’s outer membrane, which forms multimeric clusters that structure water into an ice-like array, promoting its phase change (Green and Warren, 1985; Govindarajan and Lindow, 1988). This trait has been linked to the increased susceptibility to frost damage above -5 °C of plants harboring populations of these bacteria on their leaves (Lindow *et al.*, 1982).

This particular IN ability of *P. syringae* and similar bacteria has been long suggested to possibly influence atmospheric processes (Sands *et al*., 1982; Morris *et al*., 2014). The freezing of the droplets that compose clouds (glaciation) is an important mechanism leading to precipitation (including hail formation), and is largely determined by IN particles present in suspension on the air. Due to the reduced vapor pressure over ice crystals than supercooled liquid water, frozen particles can accumulate humidity and grow to sizes large enough to start the precipitation process inside the cloud (Möhler *et al.,* 2007; Murray *et al.,* 2012). This is known as the Wegener-Bergeron-Findeisen process, and its significance is evidenced by observations such as that the ice phase of clouds is the main source of rain in continental areas across the globe (Mülmenstädt *et al.,* 2015). An additional mechanism that can amplify the influence of IN is the Hallett-Mossop process, the rapid multiplication by orders of magnitude of secondary ice crystal fragments caused by the riming and splintering of primary ice surfaces (Hallett and Mossop, 1974). Since this occurs predominantly between -3 and -8 °C, temperatures where biological IN are the major active nuclei present in the environment (Murray *et al.,* 2012), this has been proposed as another potential contribution that organisms like *P. syringae* can have for the precipitation cycle (Morris *et al.,* 2014).

In addition to cloud glaciation, microbial cells can also exhibit activity as cloud condensation nuclei (CCN) in warm clouds (Bauer *et al.,* 2003). These aerosol particles are essential for the condensation of water vapor into the liquid droplets that make up clouds. Interestingly, besides their IN activity, studies with bacteria of the *Pseudomonas* genus have also shown their ability to produce biosurfactants that can act as highly efficient CCN (Ahem *et al*, 2007; Ekström *et al.,* 2010; Renard *et al.,* 2016). Further research of microbial life in clouds also includes the effects of cells on clouds chemistry, particularly in relation to the metabolism of organic compounds in its aqueous phase (Delort *et al.,* 2017).

Multiple works have reported the presence of cultivable *P. syringae* and other ice nucleating bacteria in rain and snow samples (Morris *et al.,* 2008; Šantl-Temkiv *et al.,* 2015, for example). Stopelli *et al.* (2017) isolated IN-active *P. syringae* with selective culture media from snow collected at an altitude of 3580 m at Jungfraujoch, Switzerland. These organisms have also been isolated directly from clouds (Amato *et al.,* 2007; Joly *et al*., 2013). Members of the *Pseudomonas* genus were the most frequently identified bacterial isolates from cloud water samples collected at the puy de Dôme summit in France (at an altitude of 1465 m) between 2007 and 2010 (Vaïtilingom *et al.,* 2012). A number of *Pseudomonas* strains were also isolated from clouds and rain at the Outer Hebrides, Scotland, although those did not present IN activity (Ahern *et al*., 2007). These evidences point to the widespread distribution of these bacteria on the atmosphere and support their relationship with clouds and the precipitation cycle.

Besides *Pseudomonas*, a concentration of about 10^4^ bacterial cells per cubic meter is estimated to be found typically over land, though this number may significantly change with the altitude, weather, season, and the underlying ecosystem (Bauer *et al*., 2002; Burrows *et al*., 2009). Particles the size of bacterial cells have a relatively long residence time on the air, on the order of days, during which they have the potential to cross long distances (Burrows *et al*., 2009; Wilkinson *et al*., 2012). Effective dispersal trough this medium is, however, conditioned to cell survival as aerosols, which is a considerable challenge in this situation.

The viability of aerosolized bacteria can be severely limited by atmospheric factors such as dehydration and exposure to ultraviolet (UV) light. Even inside clouds, microorganisms can still be subjected to UV, and additionally to low temperatures, freezing, and chemical stresses such as low pH and oxidizing species (Delort *et al*., 2017). An important mean of escape from this situation may be through precipitation, which can be facilitated by the IN activity of the biological particle, as mentioned above. Both field measurements and laboratory studies have shown that cells with this activity can be preferentially precipitated from clouds in this manner, more so than non-nucleating particles (Amato *et al.,* 2015; Stopelli *et al.,* 2015; Stopelli *et al.,* 2017). In this manner, ice nucleation could be a valuable feature for airborne bacteria to return to the ground and again be able to multiply and propagate.

In this work, two strains of *P. syringae* (pv. *syringae* and pv. *garcae,* based on Gongalves and Massambani (2011)) were tested against simulated conditions that these bacteria would be exposed on the high atmosphere: UV and desiccation (which, indeed, can also be found on their natural plant surface habitat). The survival of the cells and their IN activity was quantified after the treatments to improve the understanding of their response to these factors in the environment. A strain of the model organism *Escherichia coli,* another Gram-negative gamma-proteobacterium like *Pseudomonas,* but non-ice nucleation active and not a common inhabitant of plant surfaces, was used for comparison.

## MATERIAL AND METHODS

### Strains, media, growth conditions, and survival quantification

*P. syringae* cells, from the strains IBSBF 281^T^ (= NCPPB 281, ATCC 19310; pv. *syringae*; isolated from *Syringa vulgaris*) and IBSBF 158 (pv. *garcae*; isolated *Coffea arabica,* where it causes the brown spot disease on leaves), were grown at 15 °C in L_NP_ medium (MOPS, 10.46 g L^-1^; KC1, 1.86 g L^-1^; NH_4_C1, 0.11 g L^-1^; Na_2_SO_4_, 1.42 g L^-1^; NaCl, 0.58 g L^-1^; MgCl_2_-6H_2_O, 0.20 g L^-1^; KH_2_PO_4_, 0.014 g L^-1^; CaCl_2_, 0.011 g L^-1^; FeCl_3_-6H_2_O, 0027 g L^-1^; sorbitol, 4 g L^-1^; pH 7.2) for 3 days to an OD of ∼0.5. This nutrient-limited medium and cultivation conditions were chosen to allow for maximum expression of the IN phenotype (Nemecek-Marshall *et al.,* 1993). Colony-forming units (CFU) were enumerated on Difco Nutrient Agar added with 2.5% glycerol (NAG) plates incubated in the dark at 20 °C. *E. coli* BL21 was cultivated at 37 °C on Difco LB Broth. Cultures were grown overnight to an OD of ∼5.0 in LB, and enumerated on LB agar plates incubated in the dark at 37 °C. All cell suspensions were diluted in saline solution (NaCl 0.9% w/v) prepared with ultrapure Milli-Q water (Millipore, Molsheim, France). Survival is expressed as the fraction N/N_0_, where “N” is the dilution-corrected number of UFC recovered after each treatment and “N_0_” is the number of initial UFC from before the experiments. Survival fraction values are presented as means of the replicates, with error bars denoting standard deviations.

### Quantification of ice nucleation activity

Ice nucleation activity for each sample was quantified on diluted cell suspensions placed as arrays of 32 drops of 10 μl on top of a paraffin-coated aluminum tray. This coating was previously applied as a 2% solution of paraffin in xylene, with the solvent removed by heat over a hot plate. The tray was covered with a transparent acrylic lid sealed on the borders by a ring made of EVA foam sheet and held in place by binder clips. This set was then positioned almost totally immersed in a low temperature circulating bath (Neslab LT-50, Newington, USA) filled with 96% ethanol. Temperature was monitored with a submerged mercury thermometer. From -2 °C, the bath temperature was reduced in 1 °C stages, which were held for at least 5 minutes. At each stage, the number of frozen drops was scored. IN concentration was calculated with the following equation adapted from Vali (1971), as commonly used for microbiological studies (e.g., Joly *et al.* (2013)): c(T) = [ln (N) - ln (N - N(T))] / A, where “c(T)” is the number of cumulative active IN per cell at temperature “T”, “N” is the number of drops tested, “N(T)” is the number of frozen drops at temperature “T”, and “A” is the number of cells per drop (determined by CFU counting). Measured IN activity values are presented as means of the replicates, with error bars representing standard deviations. Ice nucleation profiles across a range of temperatures for the 281 and 158 P. *syringae* strains are presented on **Figure 1.** Interestingly, the pv. *syringae* strain had a stronger measured IN activity than the pv. *garcae* strain, different to what was previously reported by Gonçalves and Massambani (2011).

**Figure 1.**
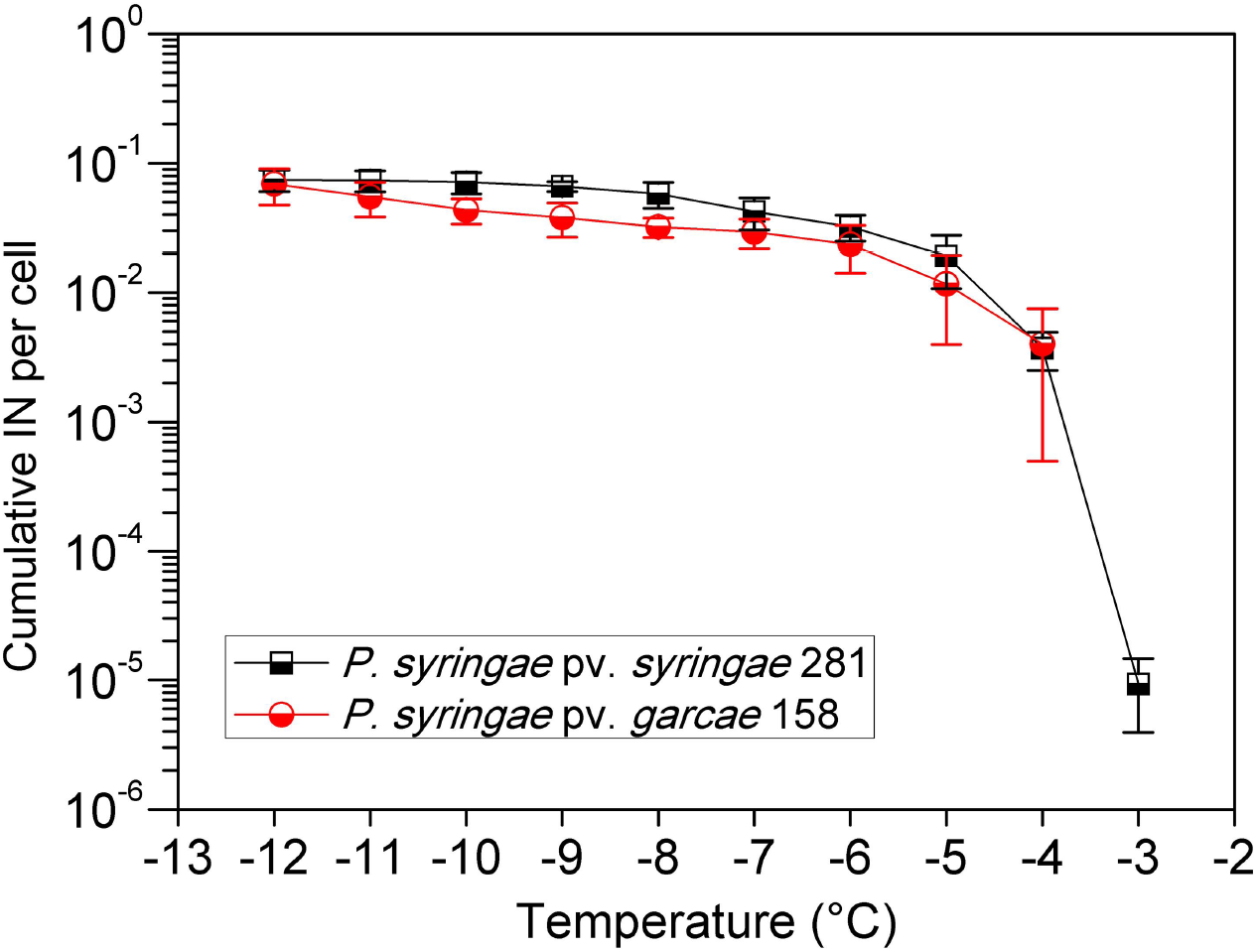
Cumulative ice nucleation spectrum of *P. syringae* pv. *syringae* 281 and *P. syringae* pv. *garcae* 158.

### UV irradiation experiments

UV-C irradiation was done with a Philips TUV-20W low-pressure mercury lamp with main emission line at 254 nm. UV-B was provided by a set of mercury lamps (two LightTech Narrow Band UV-B 20W and one Philips TL 20W/01 RS) with main emission line at 312 nm. A solar simulator (Oriel Sol-UV-2, Bozeman, USA) with a 1000 Watt xenon arc lamp was used for the “environmental UV” irradiation experiments. This source’s output covers partially the UV-B and UV-A ranges with a spectrum similar to the one found at directly Sun-exposed environments on Earth (with negligible UV-C), except most of the visible light is removed by an optical filter. Spectra of the lamps used for these experiments can be observed in **Figure 2.** Irradiation intensities and fluences were measured by a radiometer (Vilber Loumart RMX-3W, Marne-la-Vallée, France) with photocells specific to wavelength ranges centered on 254 nm at the UV-C (CX-254), 312 nm at the UV-B (CX-312), and 365 nm at the UV-A (CX-365).

**Figure 2.**
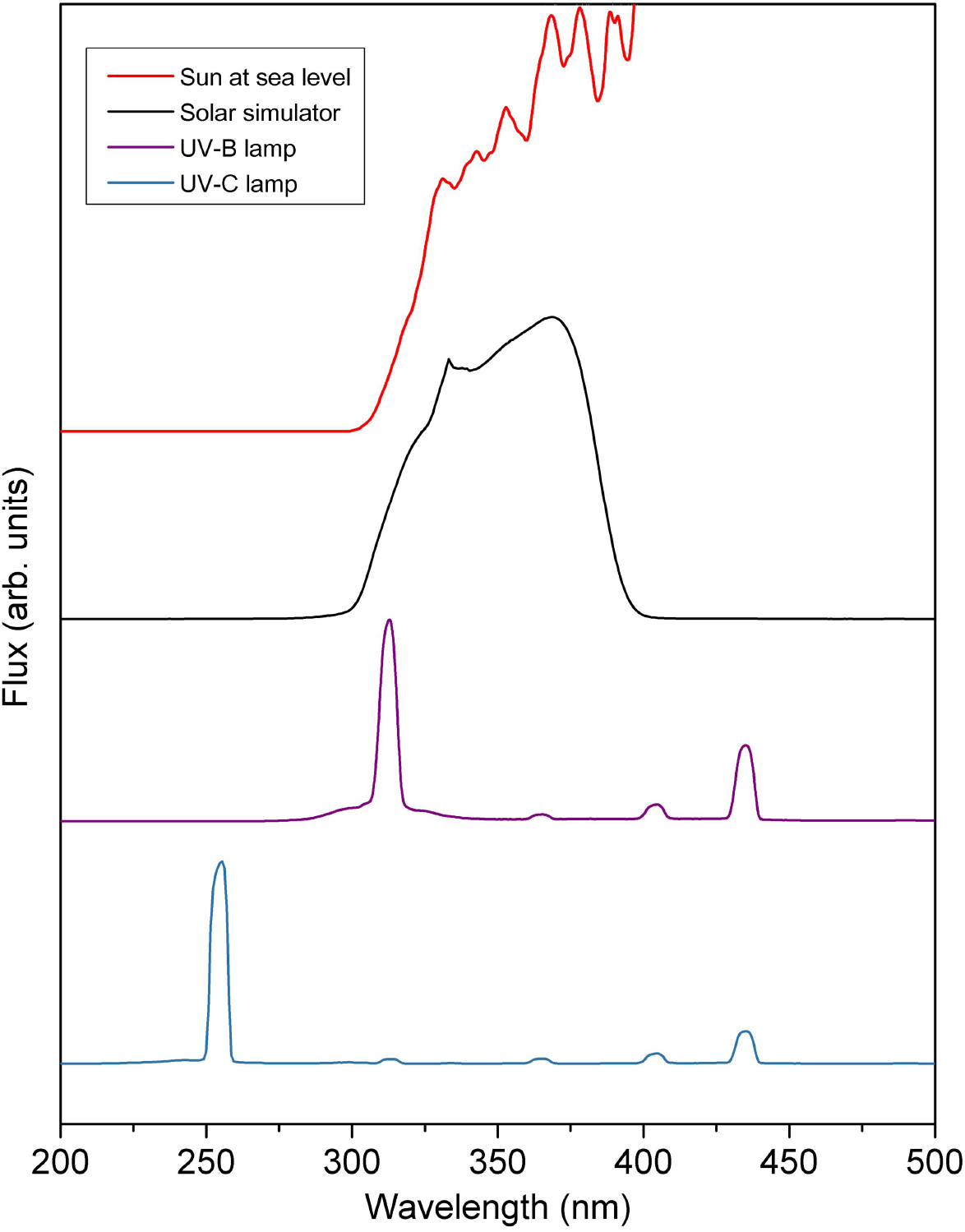
Comparison of the spectra of the lamps used for the experiments, from top to bottom (in arbitrary flux unities): of the Sun over Earth’s surface (ASTM G173-03, smoothed), of the Sol-UV-2 solar simulator, and of the UV-B and UV-C lamps, measured with an Ocean Optics QE65000 spectrometer.

For the UV-C and UV-B assays, cultures were diluted 100-fold in saline solution to a final volume of 8 ml in autoclaved 7 cm diameter glass petri dishes (without the lids). The samples were irradiated under orbital shaking while the fluences were monitored in real time by photocells placed next to the dishes. Aliquots taken at fluence intervals were diluted and plated for survival quantification by enumeration of CFU. For the “environmental” UV experiments, the samples were irradiated on top of a frozen foam block, also under shaking, so as to control their temperature to near 0 °C. This was done during these more prolonged exposures to avoid evaporation and to prevent excessive heating – which can itself affect the cells’ IN by leading to the disaggregation of the ice nucleation protein clusters on the cells’ outer membranes (Nemecek-Marshall *et al*., 1993). The intensities were measured beforehand as 75.5 W/m^2^ for the UV-A and 48.7 W/m^2^ for the UV-B, at the ranges read by the radiometer (centered on 365 and 312 nm). Aliquots from the exposed samples were then taken after determined time intervals, corresponding to UV-A and UV-B fluences which could be calculated afterwards from the intensity values.

Separate experiments were performed to test the effects of UV over the cells’ IN. Longer exposures were used in these assays since preliminary trials showed no effects of smaller fluences on IN activity. With the solar simulator, samples were exposed for 120 minutes at the same UV intensities as before, equivalent to a total 545 kJ/m^2^ of UV -A and 348 kJ/m^2^ of UV-B (at the ranges read by the radiometer), twice as much as the largest fluence of the survival tests. In this case, the diluted cultures were exposed as 2 ml volumes in 3 cm diameter dishes, allowing more samples to be placed below the source’s focus at the same time. The control samples, “0 min”, were aliquoted from the dishes before the beginning of the experiments and stored until the end of the irradiation when their IN activity was quantified parallel to the “120 min” samples.

### Desiccation experiments

Desiccation assays were performed with 10 μl volumes taken directly from the cultures and deposited in the internal wall of horizontally positioned autoclaved 1.5 ml microcentrifuge tubes. The tubes were then placed inside sealed recipients containing either silica gel beads or water-saturated MgCl_2_. These treatments were used to provide controlled relative humidity (RH) values below 5% and of about 33%, respectively (Winston and Bates, 1960). A <5% RH is typical for high altitudes, considering water vapor sources at the ground surface. Hydrated controls were prepared by adding 10 μl from the cultures to a total 1 ml in tubes with saline solution (10^-2^ dilution). All samples were stored for 6 days inside an incubator at 20 °C. During this, the temperatures were monitored with mercury thermometers and were found to remain stable. At the end of this period, the dried samples were resuspended with 1 ml saline solution (10^-2^ dilution in relation to the cultures). Along with the hydrated control cell suspensions, these tubes were diluted for survival determination by plating and for IN quantification. Survival was calculated relative to initial controls aliquoted from the cultures, plated before the beginning of each experiment.

## RESULTS

The survival curves to UV-C radiation (254 nm) of *P. syringae* pv. *syringae* 281 and P. *syringae* pv. *garcae* 158 were relatively similar to *E. coli*‘s, though 158 was slightly more tolerant **(Figure 3)**. Under our tested conditions, a fluence of 30 J/m was enough to reduce the CFU counts of 281 to about 10% of the initial population (1 log decrease), compared to 50 J/m for an equivalent reduction in 158.

**Figure 3.**
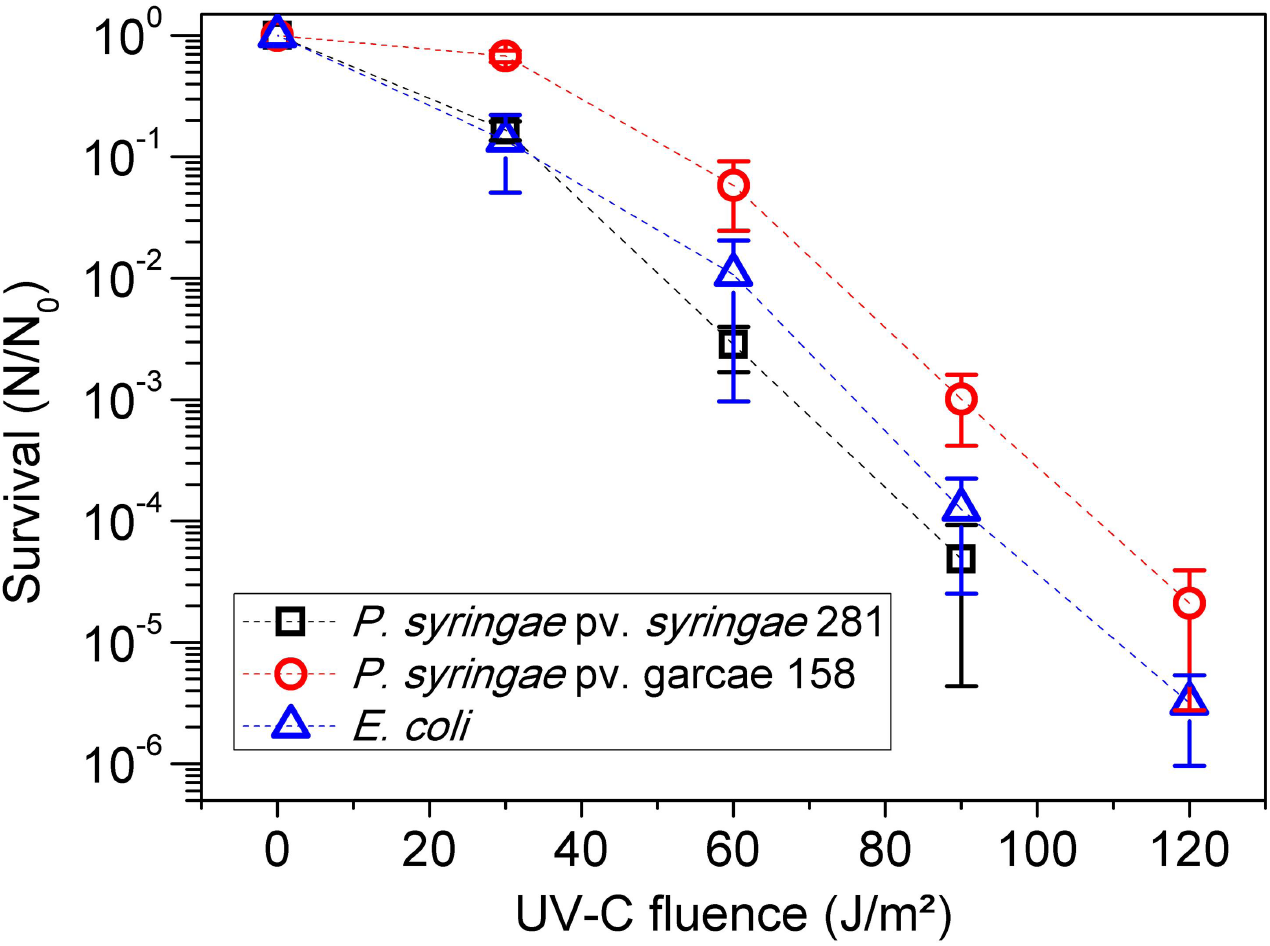
Survival curve to UV-C (254 nm) radiation of *P. syringae* pv. *syringae* 281, P. *syringae* pv. *garcae* 158, and *E. coli*.

For the UV-B (312 nm) assays, the observed survival of the *P. syringae* strains was considerably greater than *E. coli* **(Figure 4)**. At a fluence of 5000 J/m, *E. coli* was inactivated by nearly 3 logs, while both 281 and 158 lost less than 1 log of viability. Treatment with higher fluences (up to 20000 J/m^2^) evidenced again a larger UV tolerance of 158 compared to 281.

**Figure 4.**
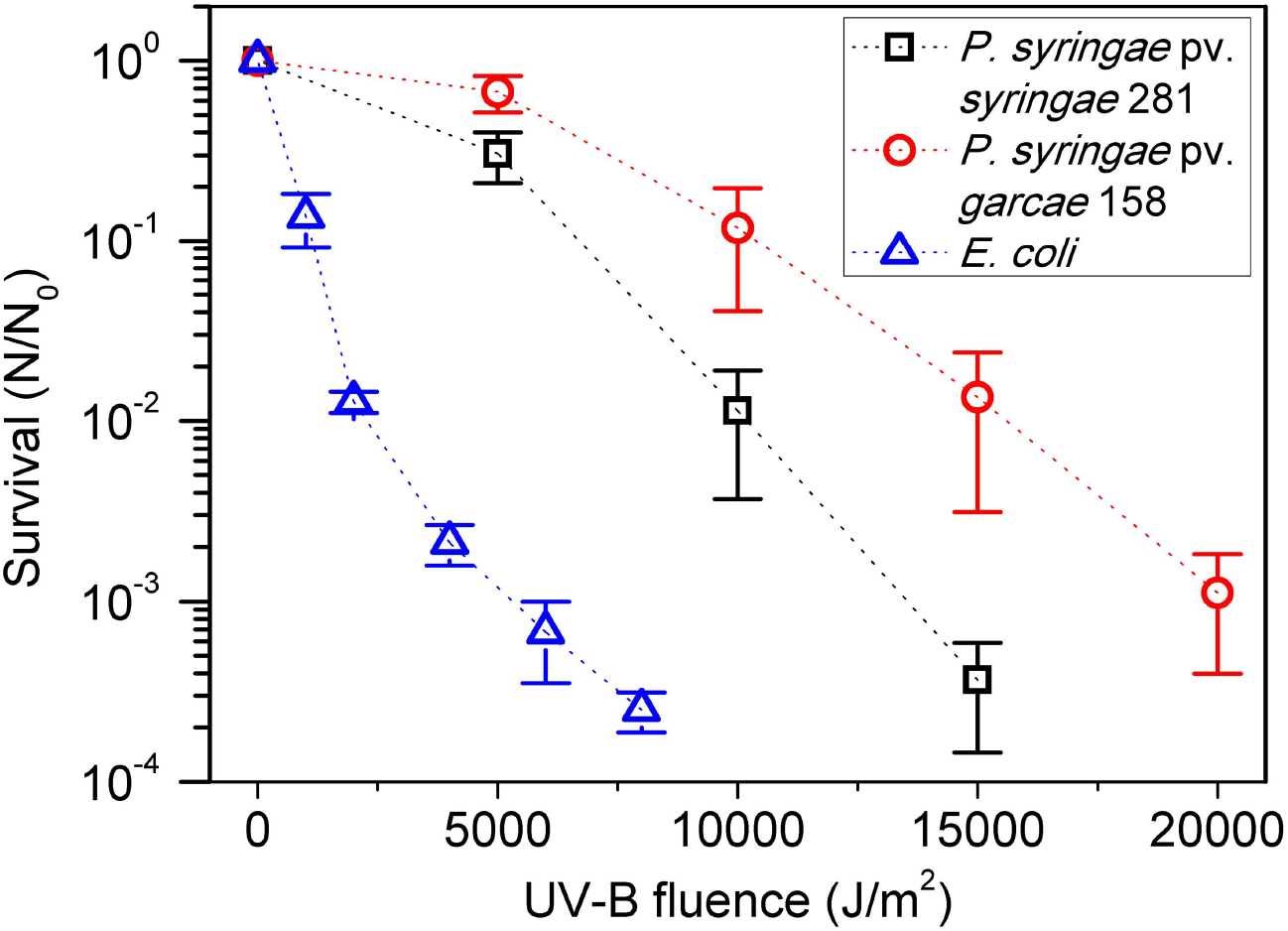
Survival curve to UV-B (312 nm) radiation of *P. syringae* pv. *syringae* 281, P. *syringae* pv. *garcae* 158, and *E. coli*.

Like the UV-B experiments, the *Pseudomonas* strains were also significantly more tolerant to irradiation with the more environmentally relevant UV range of a solar simulator when compared to *E. coli* **(Figure 5)**. A 60 minute exposure, at the intensities used for the assays, did not reduce by more than 1 log the viability of 281 (32±16% survival) or 158 (25±14% survival), while *E. coli* survived at only 0.2±0.1%. Interestingly, the response of both *P. syringae* strains was much more similar to the tested UV-A + UV-B fluences used for these experiments, where both curves were nearly overlapping, than for the monochromatic UV-B.

**Figure 5.**
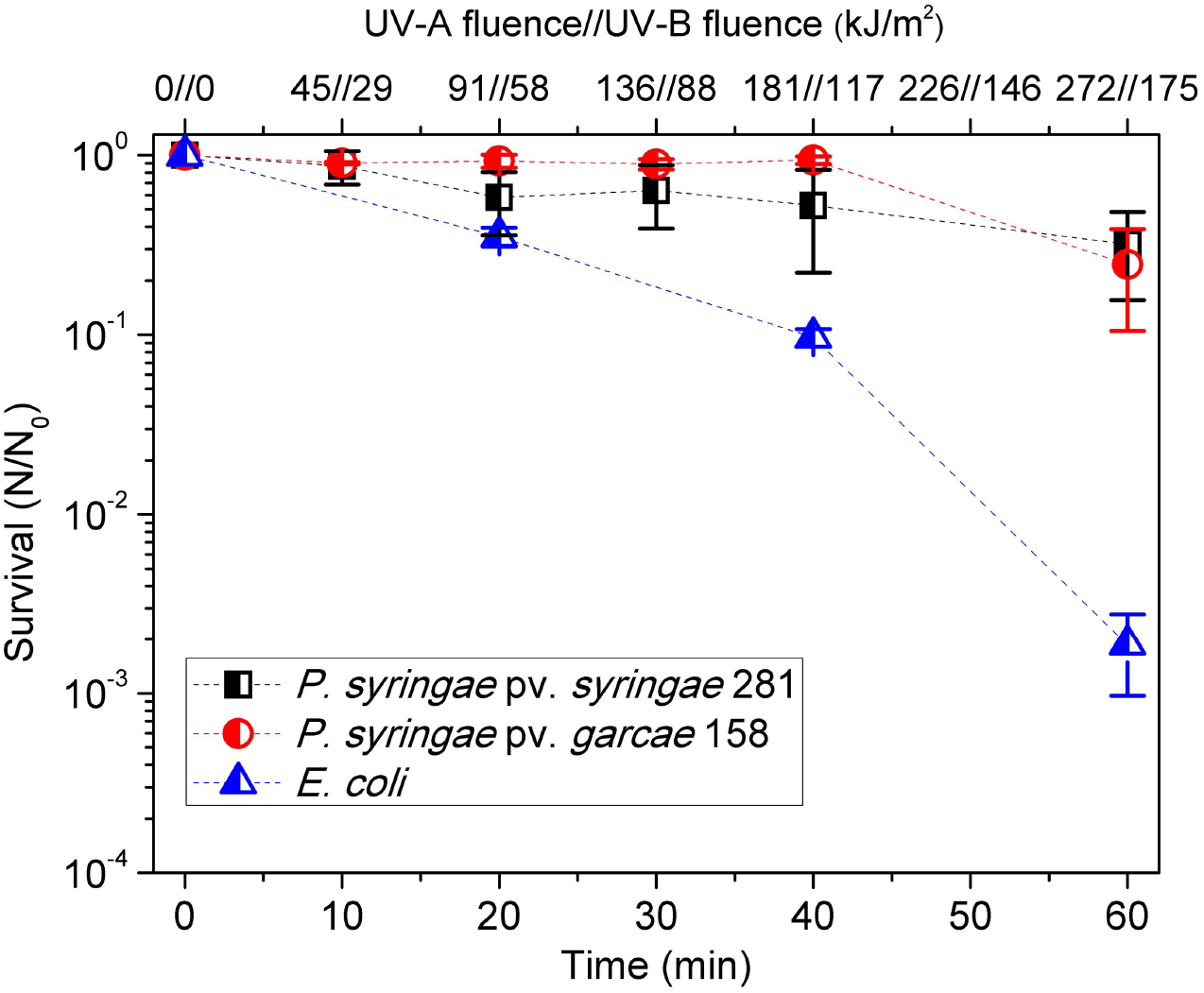
Survival curve to “environmental” TJV radiation (Uy-A + UV-B) of *P. svringae* pv. *svringae* 281. *P. svringae* pv. *garcae* 158. and *E. coli*.

The IN activity of the cells at -5 °C was measured after irradiation for 120 minutes of simulated “environmental” UV **(Figure 6)**. While 281 maintained its initial IN concentration, 158 presented an up to 10-fold decrease from the typical 10^-2^ -10^-1^ nuclei per cell of this strain at this temperature.

**Figure 6.**
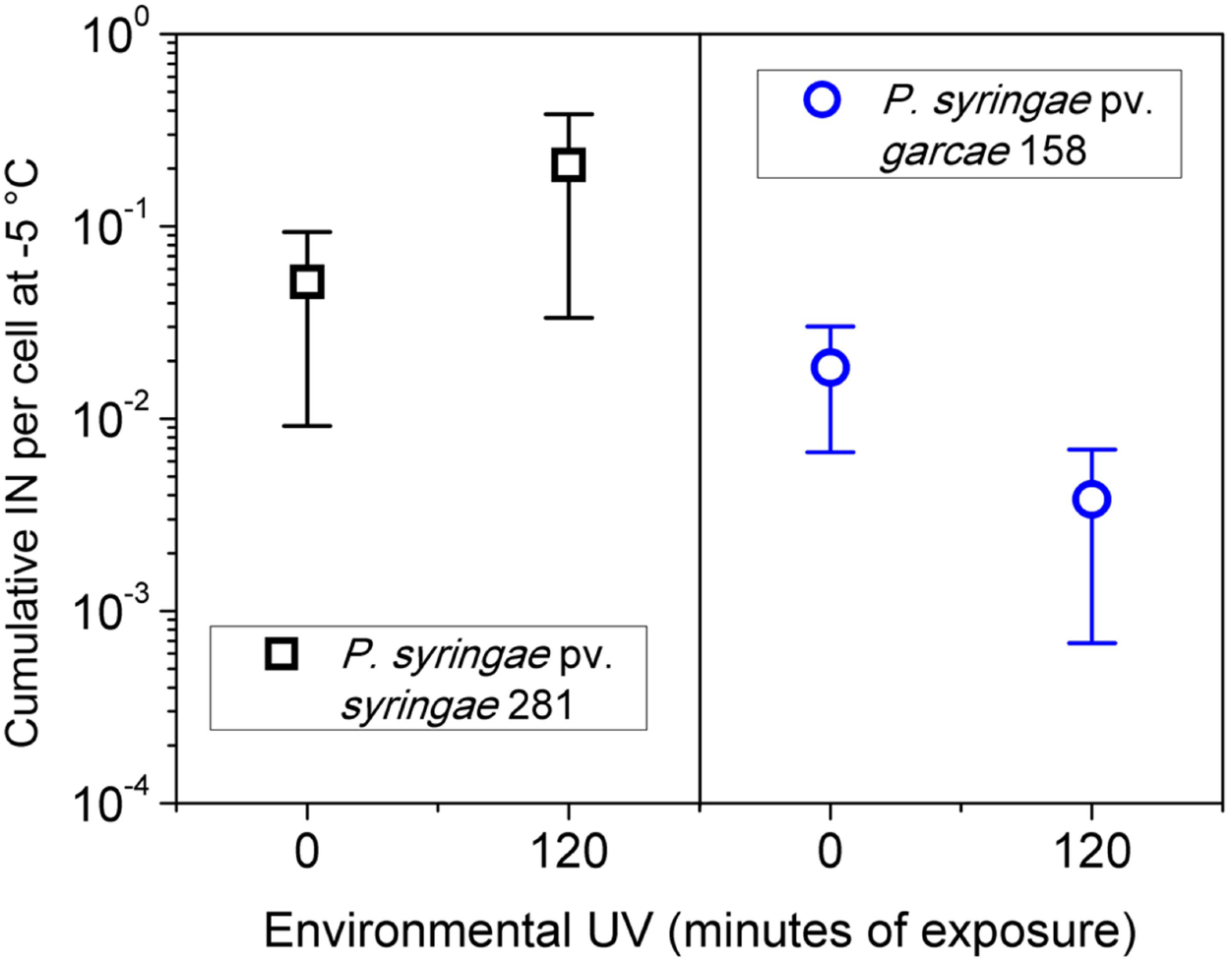
Concentration of cumulative ice nuclei per cell at -5 °C of *P. syringae* pv. *syringae* 281 and *P. syringae* pv. *garcae* 158 samples exposed for two hours to “environmental” UV radiation (UV-A + UV-B), compared to non-irradiated controls (“0 min”).

For the desiccation assays, cells were kept for 6 days inside recipients with controlled RH at 20 °C. Those results are presented on **Figure 7**. The largest tolerance was exhibited by the 158 strain at RH <5%, in which 22±8% of its initial population survived. At the same treatment, the viability of 281 was reduced by 3 to 4 logs. At RH 33%, the surviving percentage of 158 was 4±2%, about 10 times more than 281. For both tested RH (33% and <5%), *E. coli* mean survival was around 3 to 4 %. Control samples, kept hydrated in saline solution during the 6-day period, remained mostly at the same initial CFU concentration, except for 281 for which the number of cells increased slightly.

**Figure 7.**
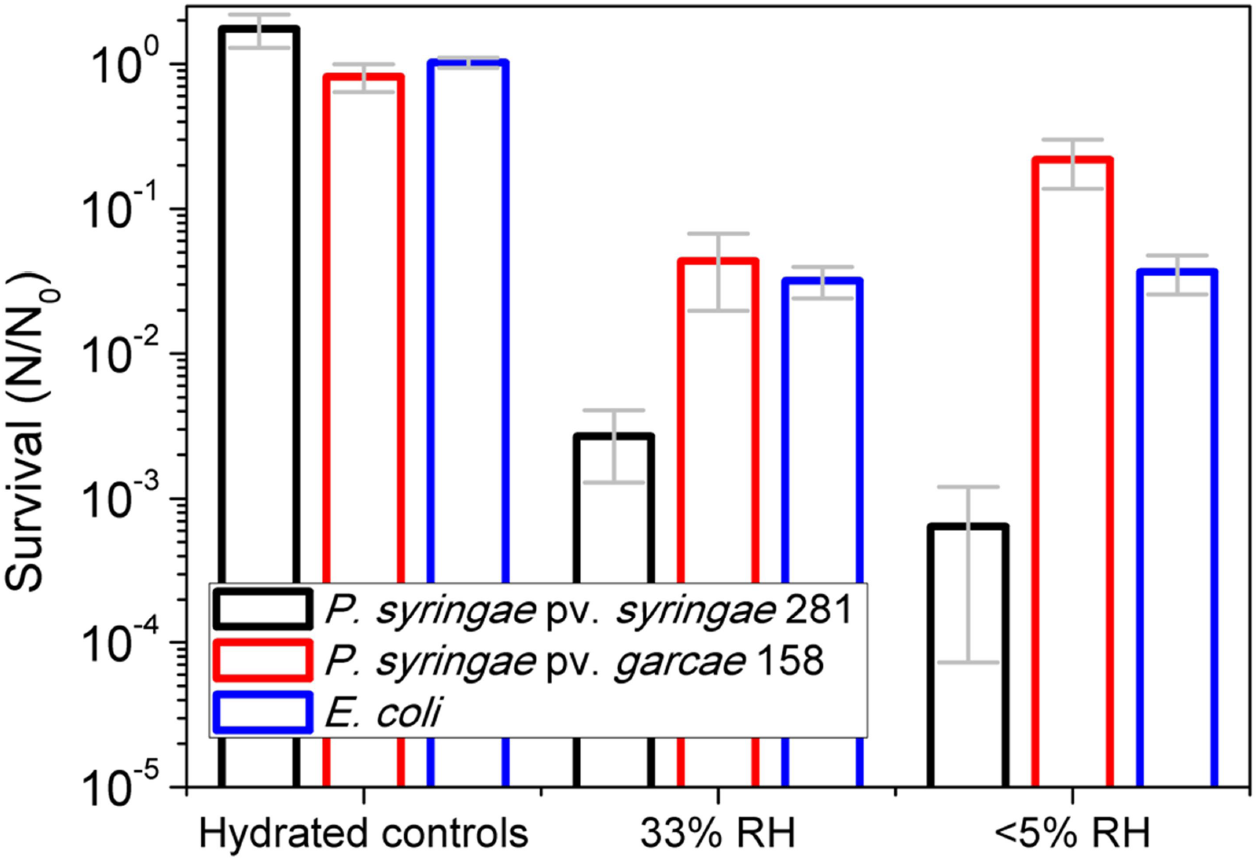
Survival of hydrated controls and samples desiccated at RH 33% and <5% for 6 days at 20 °C of *P. syringae* pv. *syringae* 281, *P. syringae* pv. *garcae* 158, and *E. coli.*

The IN activity of the desiccated *P. syringae* strains was strikingly different **(Figure 8)**. The 281 strain presented a mean cumulative nuclei concentration of about 3-4×10^-3^ per cell at -5 °C after being resuspended from the RH 33% and RH <5% treatments. Its hydrated controls remained at typical values for these cultures, at 1.1±1.0×10^-1^. However, the IN concentrations for the 158 strain, which are normally similar to those of 281, were lower by 10^3^-10^4^ times, even for hydrated cells. Its mean measured activities (concentration per cell at -5 °C) were from 1×10^-4^ to 9×10^-6^, after the 6-day period at 20 °C.

**Figure 8.**
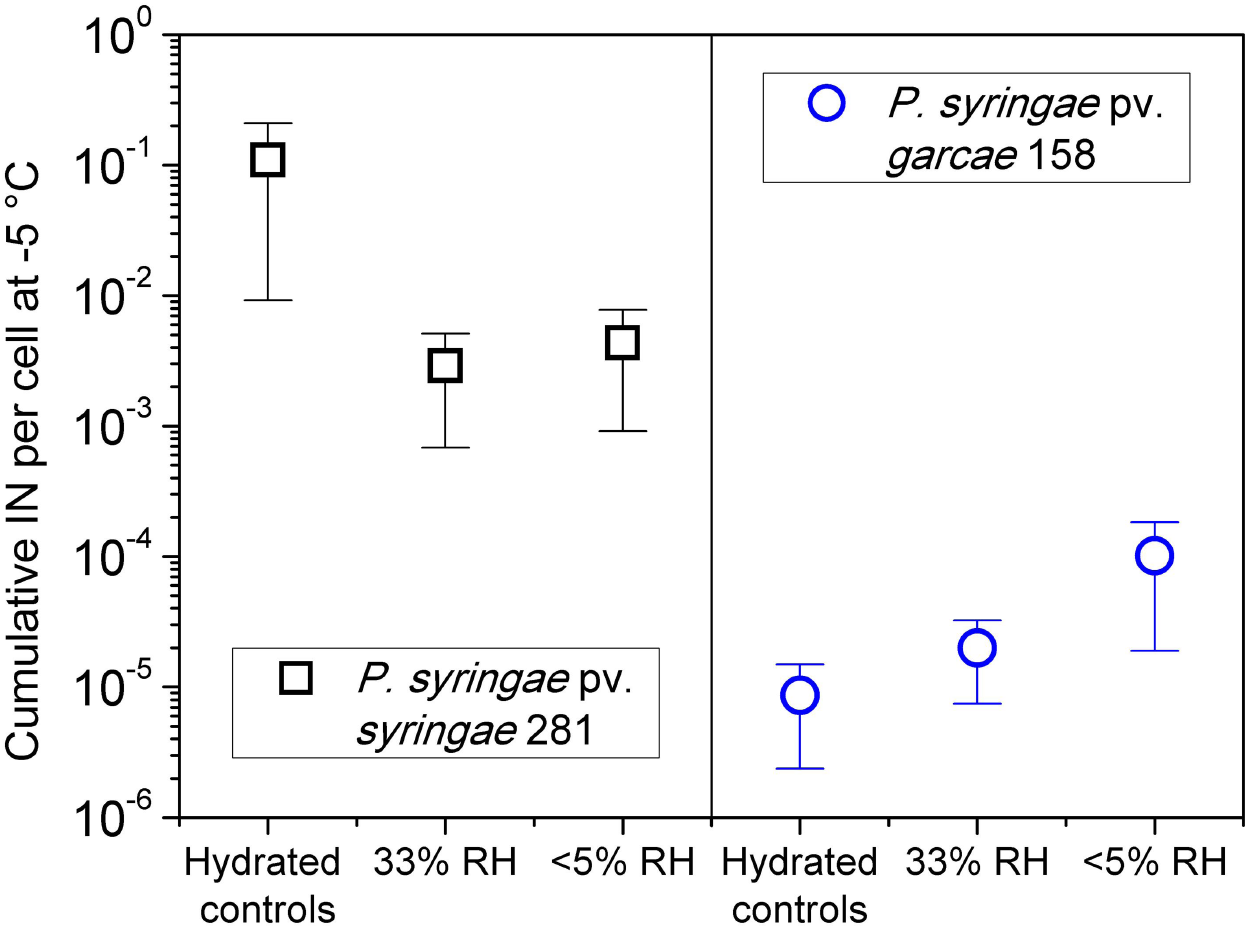
Concentration of cumulative ice nuclei per cell at -5 °C of *P. syringae* pv. *syringae* 281 and P. *syringae* pv. *garcae* 158 samples desiccated at RH 33% and <5% for 6 days at 20 °C and their respective hydrated controls.

## DISCUSSION

### P. syringae survival after exposure to different UV ranges

Both *P. syringae* strains were found to be very sensitive to UV-C, exhibiting survival comparable to the non-UV tolerant bacterium *E. coli* **(Figure 3)**. Solar radiation emitted at this range (<280 nm) does not reach Earth’s lower atmosphere (the troposphere), being completely absorbed at the stratosphere. Despite that, the 254 nm wavelength produced by low-pressure mercury lamps is widely used to assess the effects of UV on bacteria in laboratory studies. It efficiently inactivates microorganisms, and, as such, is commonly referred to as “germicidal UV” (Coohill and Sagripanti, 2008). Nevertheless, some bacteria, such as *Deinococcus radiodurans*, show extreme tolerance to UV-C radiation (Slade and Radman, 2011). For comparison, a UV-C fluence of about 1000 J/m^2^ is required to inactivate 90% of its population (Pulschen *et al*., 2015), while the highest fluence tested in this work was 120 J/m^2^ **(Figure 3)**.

UV-C causes the photochemical formation of dimers between adjacent pyrimidine bases on the cells’ DNA. The main photoproducts generated are cyclobutane pyrimidine dimers (CPDs) and (6-4) pyrimidine-pyrimidone photoproducts (6-4 PPs), which block transcription and DNA replication with potential lethal effects for the cell (Coohill and Sagripanti, 2008). A widespread UV resistance mechanism is photorepair, where a specific enzyme, called photolyase, binds pyrimidine dimers in the cell’s DNA and uses luminous energy (UV-A or visible light) to independently repair the damage. Otherwise, cells may rely on the nucleotide excision repair (NER) pathway, a multi-step (and more energetically demanding) enzymatic process that removes a patch of the lesioned strand that is subsequently resynthesized (Coohill and Sagripanti, 2008; Slade and Radman, 2011; Meador *et al.,* 2014). The organisms tested in this work all probably possess these very common repair pathways, including other relevant systems such as base excision repair (BER) and homologous recombination, which have already been characterized in *E. coli* and can be identified in *P. syringae* genomes (Feil *et al.,* 2005).

In addition to DNA repair, it was found that protection of cellular proteins from reactive oxygen species (ROS) generated upon exposure to UV-C is an important factor for the survival of *D. radiodurans* to this condition (Krisko and Radman, 2010; Slade and Radman, 2011). This was compared to *E. coli,* which is UV-sensitive and whose proteome is severely oxidized by UV-C. Its damaged repair machinery is prevented from correcting DNA injuries and lead, ultimately, to cell death (Krisko and Radman, 2010). In this manner, avoidance of ROS formation and effective quenching of these species represent another important UV tolerance mechanism. Still, different UV wavelengths induce distinct biological effects (Santos *et al.*, 2013), which may require specific adaptations to allow the survival of the irradiated organism.

Covering the wavelengths between 280 and 320 nm, UV-B is the most energetic, and potentially damaging, range of solar radiation that reaches the ground, despite most of it being absorbed by the ozone layer. At higher elevations, the UV flux is increased in relation to lower altitudes, more so at the UV-B region (Blumthaler *et al.*, 1992). In agreement to that, Wang *et al.* (2014) measured a more severe DNA damage in plants at an altitude of 1700 m than at 300 m. UV-B is known to directly create DNA photoproducts, like UV-C, but is also capable of causing significant oxidative damage from ROS (Santos *et al.*, 2013). In contrast to the UV-C assays, the *P. syringae* strains were significantly more resistant to this UV range (at a wavelength of 312 nm) than *E. coli* **(Figure 4)**. However, it must be recognized that all these tested organisms are still much more sensitive than *D. radiodurans*, for which over 120000 J/m of UV-B is required to inactivate about 90% of its population at identical experimental conditions (Pulschen *et al.*, 2015).

The strain 158 (*P. syringae* pv *garcae*) was distinctly more tolerant than 281 (*P. syringae* pv. *syringae*) to the UV-B treatment, more perceptible at the higher fluences tested. This evidences substantial differences between the strains (which can also be observed on the results of the other experiments discussed further below), despite both being classified as the same species. In fact, *P. syringae* pv. *garcae* can be distinguished as belonging to a separate, discrete genomospecies than *P. syringae* pv. *syringae* on the basis of DNA differences, though a lack of discerning phenotypic characteristics has prevented its reclassification with other strains within the “*P.coronafaciens*” taxon (Gardan *et al.*, 1999).

Some *P. syringae* strains possess the error-prone DNA polymerase V encoded by the *rulAB* operon, which is responsible for translesion synthesis over damaged DNA template strands. The expression of this polymerase is induced by UV-B and confers increased resistance towards irradiation at the cost of increased mutability (Kim and Sundin, 2000). This operon is most commonly found in plasmids and its occurrence is variable within the species, even in strains of the same pathovar (Sundin and Murillo, 1999; Feil *et al.*, 2005). Its presence in the *Pseudomonas* strains tested in this work would thus have to be individually verified if this tolerance factor was to be attributed to them.

The UV-C and UV-B lamps used in this study are sources of narrow band radiation, much different from the continuous spectrum found in the environment **(Figure 2)**. For a more accurate representation of the environmental UV, a solar simulator emitting UV-A and UV-B was used. Under these conditions, the *P. syringae* strains were much more resistant than *E. coli*, surviving about 2 logs more at the 60 minutes exposition **(Figure 5)**. This can be partially seen as a consequence of the observed higher tolerance of *Pseudomonas* to the UV-B **(Figure 4)**, though the presence of UV-A (320 − 400 nm) can significantly affect some organisms under this type of irradiation. For example, *D. radiodurans* is surprisingly sensitive to this higher wavelength fraction of the environmental UV (Slade and Radman, 2011; Pulschen *et al.,* 2015).

The deleterious biological effects of the UV-A range are mostly linked to ROS production, damaging, albeit indirectly, the cells’ DNA, protein, and lipids (Santos *et al*., 2013). Even so, though much less efficiently than UV-B, UV-A is also able to form CPDs, and it can additionally cause the photoisomerization of 6-4 PPs into its Dewar valence isomers, another type of environmentally-relevant DNA damage (Meador *et al.*, 2014). It was reported that the alternative sigma factor RpoS is an important element in the survival of *P. syringae* pv. *syringae* under natural sunlight (Miller *et al.*, 2001). Genes regulated by this protein have already been characterized in *E. coli* as involved in the cellular response to oxidative stress, including DNA repair and ROS quenching functions. Inactivation of *rpoS* lead to increased sensibility of *P. syringae* to solar UV, evidencing its important role in UV tolerance for this organism (Miller *et al.*, 2001).

Joly *et al.* (2015) exposed two *P. syringae* isolates collected from cloud water for 10 hours to total final fluences of 85.7 kJ/m^2^ of UV-A and 27 kJ/m^2^ of UV-B at 5 °C. After this period, the isolates suffered virtually no viability loss. This is in agreement to the results presented on **Figure 5**, where the *Pseudomonas* strains are shown to tolerate acute expositions to considerably larger fluences than the ones used for this previous study. Still, these authors used a fluorescent lamp with a much different spectrum from the solar simulator, including a large amount of visible light. Attard *et al.* (2012) used a more similar radiation source, a 1000 Watt xenon lamp, at an total UV-A intensity of 33 W/m. Interestingly, exposure for 42 hours, in distilled water at 17 °C, only reduced the viability of three different *P. syringae* strains by about 90% in relation to non-irradiated controls. UV-B measurements (if significant) were not provided, but the final calculated UV-A exposure was of almost 5000 kJ/m^2^. Possibly, this less acute UV exposition, at a lower intensity, could have improved the tolerance of the strains.

It is important to note that, due to differences between the UV-B lamps’ line emission at 312 nm and the solar simulator broad spectrum (∼290-400 nm, **Figure 2**), the biological response to both sources is not exactly equivalent. Thus, the survival curves presented for UV-B and “environmental” UV cannot be directly compared on the basis of the measured fluences, considering the radiometer’s probes read a spectral range, not a single wavelength.

Another relevant difference of the simulated environmental UV from the other assays is the possibility that the organisms may have been able to perform photorepair during the irradiations. For the UV-C and UV-B experiments, cells were exposed for a few minutes and then incubated in the dark after plating. Exposure to the solar simulator, instead, was performed for extended periods and in the presence of UV-A, which is required for the activity of the photolyase enzyme. In this manner, the cells may have been able to repair while being irradiated at least part of the received DNA damage (in the form of pyrimidine dimers from the UV-B). It is also worth remembering that the “environmental” UV experiments were carried out at low temperatures, similar to Joly *et al.* (2015), above a frozen foam block. This was done to prevent sample heating and excessive evaporation of the cell suspension during the prolonged irradiation. Additionally, this may better mimic the conditions of bacteria in cloud water at high elevations where lower temperatures prevail.

In relation to the UV-B intensity used for the “environmental” UV assays, 48.7 W/m^2^, it is equivalent to about 5.2 times the value at an altitude of 850 m in São Paulo, Brazil (9.3 W/m^2^) and 3.1 times the value at an altitude of 5091 m at the Atacama Desert, Chile (15.6 W/m^2^), as reported in a previous paper (Pulschen *et al*., 2015). Those measurements were performed by the same probes used for the solar simulator with a Vilber Loumart radiometer, under clear sky conditions, at noon, during summer, and at similar latitudes. Taking the UV-B as the most biologically relevant radiation range for comparison, *P. syringae* strains can be expected to tolerate hours of direct sunlight exposure, even without attenuating factors such as cloud coverage and association to cell clusters, mineral particles or organic fragments. Of course, those factors can become increasingly important for survival at higher altitudes and for longer periods.

To test if the *Pseudomonas* culture conditions somehow favored the greater survival of these cells under the “environmental” UV, *E. coli* cultivated in L_NP_ was also tested. Interestingly, its survival was even inferior to LB-grown cells after one hour of irradiation **(Figure S1)**. In this manner, a poorer growth medium does not seem to contribute to UV tolerance.

### Ice nucleation activity following UV irradiation

The IN activity of the cells was quantified for cells irradiated for two hours under the solar simulator, twice as long as the survival tests. Both *P. syringae* strains typically exhibited a concentration of cumulative ice nuclei per cell of 10^-2^ -10^-1^ at -5 °C **(Figure 1)**, which was also seen for the control samples (“0 min”) for this UV assay **(Figure 6)**‥ After the exposure, 281 cells seemed to maintain it’s measured IN activity at this range, with an even larger mean value. However, 158 suffered a decrease of up to 1 log at this temperature, though the mean IN concentration value was reduced by only about 5 times.

This reduction was still well above the limit of detection of the experiments, which was around 6×10^-4^-3×10^-4^ nuclei per cell at the tested conditions (32 drops of 10 μl from a 10^-5^ dilution of a culture with 5×10^8^ -1×10^9^ cells/ml). Smaller dilutions, though, would equal lower detection limits (10 times lower if a 10^-4^ dilution from the culture was used, for example). For the desiccation assays described below, higher cell concentrations had to be used to enable detection of fewer nuclei.

The three different strains of *P. syringae* irradiated by Attard *et al.* (2012) (33 W/m^2^ of UV-A for 42 hours in distilled water at 17 °C, which led to a reduction of about 90% in viability) presented either a non-significant difference in IN activity at -5 °C or suffered a relatively small 10-fold reduction. These authors discuss how dead bacteria could maintain their IN activity as long as cell integrity is not disrupted, preserving the large ice nucleation protein aggregates on the cells’ outer membranes. In fact, experiments with very large fluences of UV-C (10000 J/m^2^), far beyond the point where no surviving cell could be expected, also yielded no difference in 281’s IN at -5 °C, and a reduction by only about 20 times for 158 **(Figure S2)**.

## Survival and ice nucleation activity of desiccated cells

Strain 281 was relatively sensitive to desiccation at both tested RH (<5% and 33%), being inactivated by over 2 orders of magnitude **(Figure 7)**. It survived in smaller numbers than LB-grown *E. coli*, which itself kept a viability of about 3 to 4% after those treatments. *E. coli* grown on L_NP_ minimal medium presented a far reduced survival, with a decrease of over 3 orders of magnitude **(Figure S1)**. Instead, 158 presented similar survival to LB-grown *E. coli* at an RH of 33%, and even larger at an RH below 5%, maintaining around one-fifth (mean value) of its initial population.

Dehydration of cells causes membrane damage, DNA strand breakage, and an increased formation of ROS from the cellular metabolism leading to protein oxidation (Mattimore and Battista, 1996; Fredrickson *et al*., 2008). Again, like for the UV assays, DNA repair and ROS avoidance should be valuable features for bacterial survival under this condition. In the environment, tolerance to desiccation could possibly be achieved by activation of the aerosolized bacteria as cloud condensation nuclei (CCN), enabling hydration of the cells from water vapor harnessed from the air. Since *Pseudomonas* were shown to be potential efficient CCN due to biosurfactant production (Ahern *et al.,* 2007; Ekström *et al*., 2010; Renard *et al*., 2016), this could be another survival mechanism at the disposition of these cells.

Despite its low tolerance to desiccation, the IN activity of 281 was relatively well preserved after this type of treatment (6 days at low RH at 20 °C, **Figure 8**). This measured decrease by roughly 30 times in IN concentration can possibly be attributed to cell membrane disruption during dehydration. Its hydrated controls seemed to maintain the full typical IN concentration per cell of this strain. In contrast, the IN activity of the more desiccation tolerant strain 158 was unexpectedly reduced even in the hydrated controls. The RH <5% samples were, in fact, slightly more active than the other treatments. The large aggregates of InaZ proteins that form at the cells outer membranes, which are essential for forming efficient IN active at relatively high temperatures such as -5 °C, are known to be particularly heat sensitive (Nemecek-Marshall *et al*., 1993). Even though the temperature of 20 °C at which the hydrated and desiccated cells were kept at during these experiments were not expected to be detrimental to biological IN (Nemecek-Marshall *et al*., 1993), further tests with new cultures of 158 were performed at 4 °C. Survival was very similar at these conditions, only the RH 33% tolerance was slightly higher **(Figure 7, Figure S3)**. These refrigerator-stored samples exhibited a significantly increased maintenance of IN at the hydrated control and RH 33% samples, while the RH <5% treatment remained mostly identical **(Figure 8, Figure S3)**. Nevertheless, these values were still lower than the typical IN concentration for this strain, signifying some other mechanism contributes to the instability of the nuclei in this case where the cells are not actively growing and their metabolism, including protein synthesis, is probably reduced.

## CONCLUSION

Bacteria potentially relevant to atmospheric phenomena due to their IN activity, *P. syringae* pv. *syringae* (strain 281) and P. *syringae* pv. *garcae* (strain 158), were exposed to laboratory simulations of individual conditions that aerosolized cells may face during aerial transport up to the clouds and high altitudes. The strains were relatively sensitive to irradiation by UV-C and UV-B lamps, but survived in substantial numbers exposure the UV-A + UV-B spectrum of a solar simulator for durations that may be equivalent to several hours in the environment. Ice nucleation activity at -5 °C of cell suspensions exposed to this treatment was – at least partially for the pv. *garcae* – maintained. Thus, it can be concluded that while solar radiation can be a serious limitation to the dispersal of *P. syringae* through the atmosphere, these bacteria are adapted to endure periods of complete exposure to sunlight, and that a relatively large subset of its population can remain capable of influencing cloud nucleation.

Desiccation is another major challenge that these bacteria may face on the environment. The response of the different strains to this stress varied substantially in the experiments, considering both survival and IN maintenance. Perhaps most interestingly, as observed for the pv. *garcae,* not even hydration can aid in the preservation of the IN of certain strains not actively growing for prolonged periods of time. Possibly, this could cause a decreased probability of being scavenged by precipitation for certain IN-producing bacteria suspended for too long on the atmosphere. For strains like the pv. *syringae* tested, this should be less of a problem.

## ACKNOWLEDGEMENTS

This work was sponsored by CAPES, FAPESP (Project 2012/18936-0), the Microsfera project (407816/2013-5) of the Brazilian Antarctic Program (ProAntar), the Brazilian National Council of Technological and Scientific Development (CNPq 424367/2016-5), INCT-Criosfera (CNPq 028306/2009), and the University of São Paulo, through the Brazilian Research Unity in Astrobiology – NAP/Astrobio. The authors would also like to thank Professor Josef Wilhelm Baader (IQ - USP) for lending the low temperature bath with which the ice nucleation experiments were performed.

---

**figure S1.**
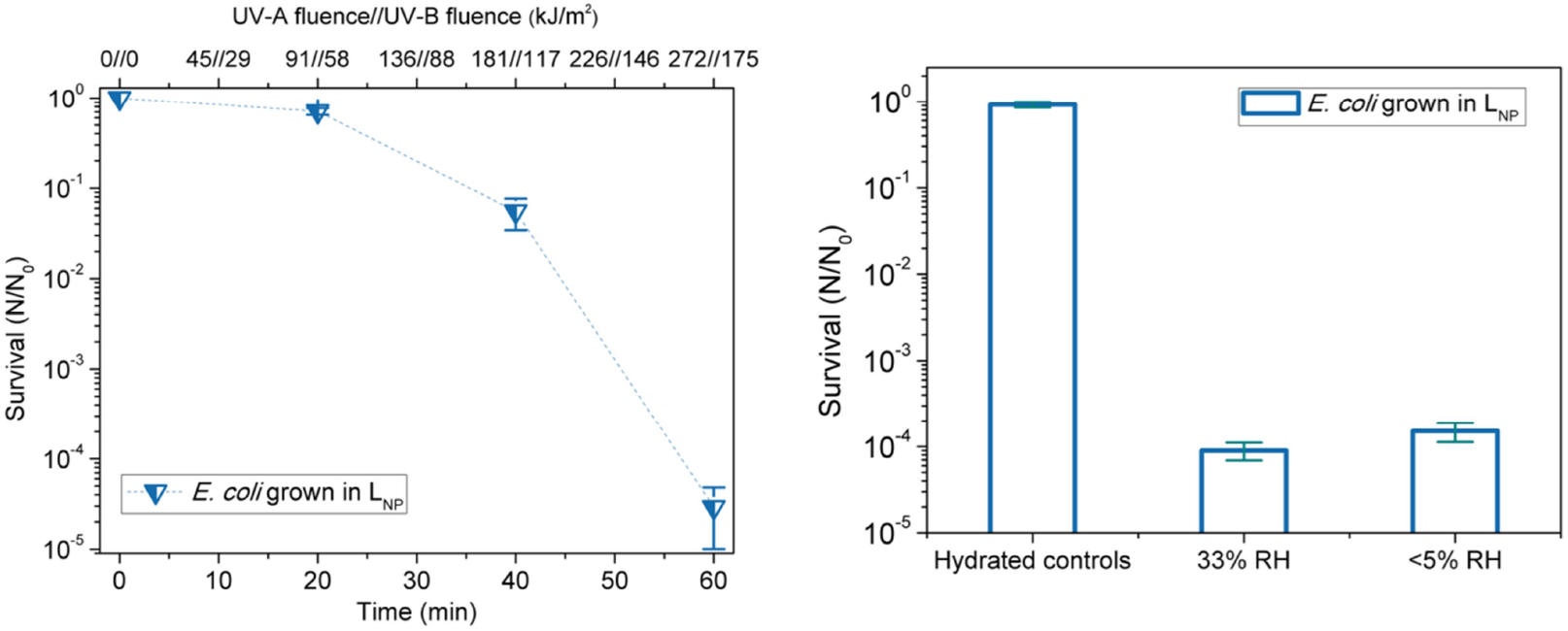
Survival of *E.coli* grown in L_NP_ medium (overnight at 37°C to OD ∼0.5) to “environmental” UV radiation (UV-A + UV-B) and to desiccation at RH 33% and <5% for 6 days at 20 °C, including its hydrated controls. Note the change in scale in the UV graph relative to **Figure 5.**

**figure S2.**
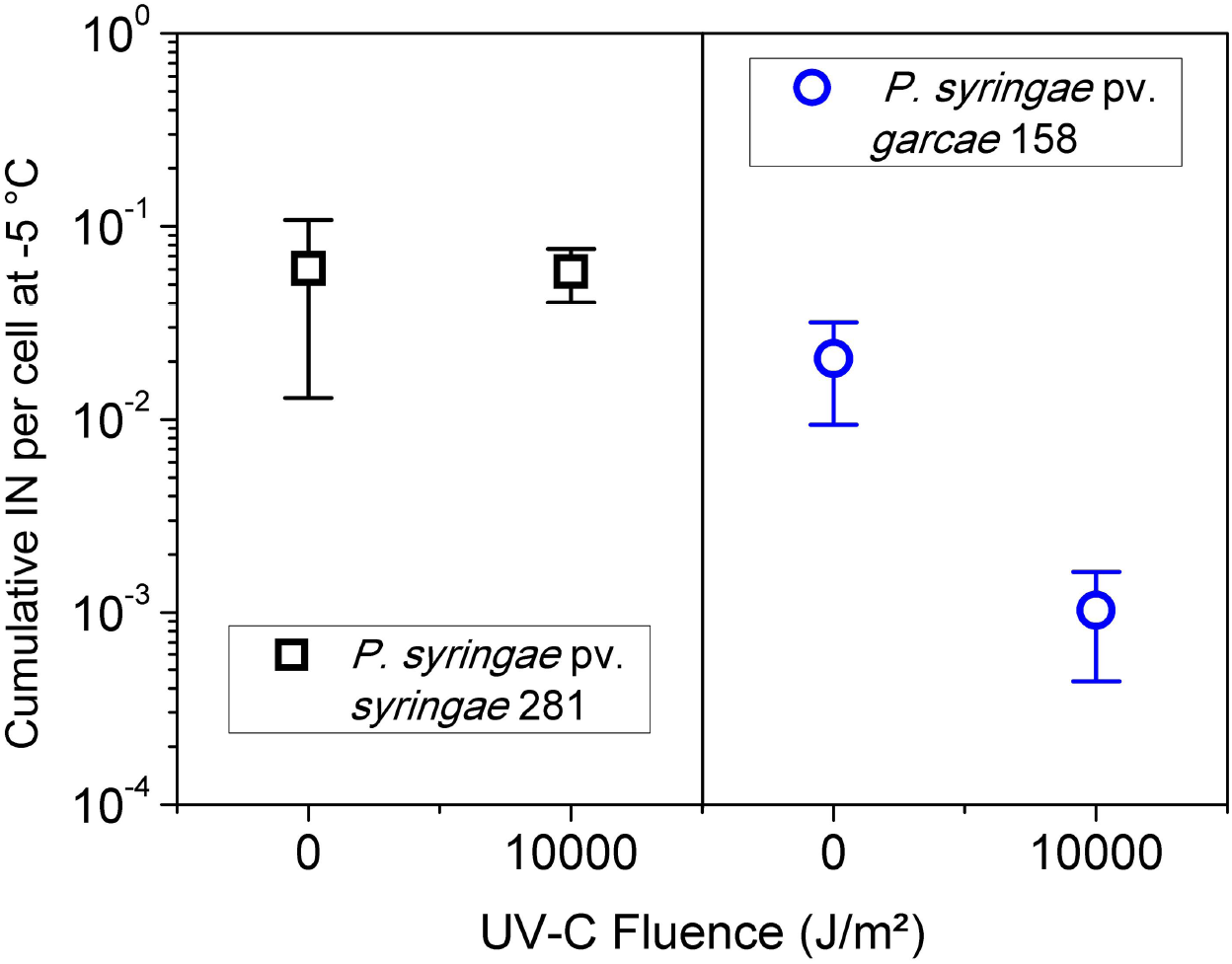
Concentration of cumulative ice nuclei per cell at -5 °C of *P. syringae* pv. *syringae* 281 and P. *syringae* pv. *garcae* 158 samples exposed to 10000 J/m^2^ of UV-C (254 nm), compared to non-irradiated controls (“0 J/m^2^”).

**figure S3.**
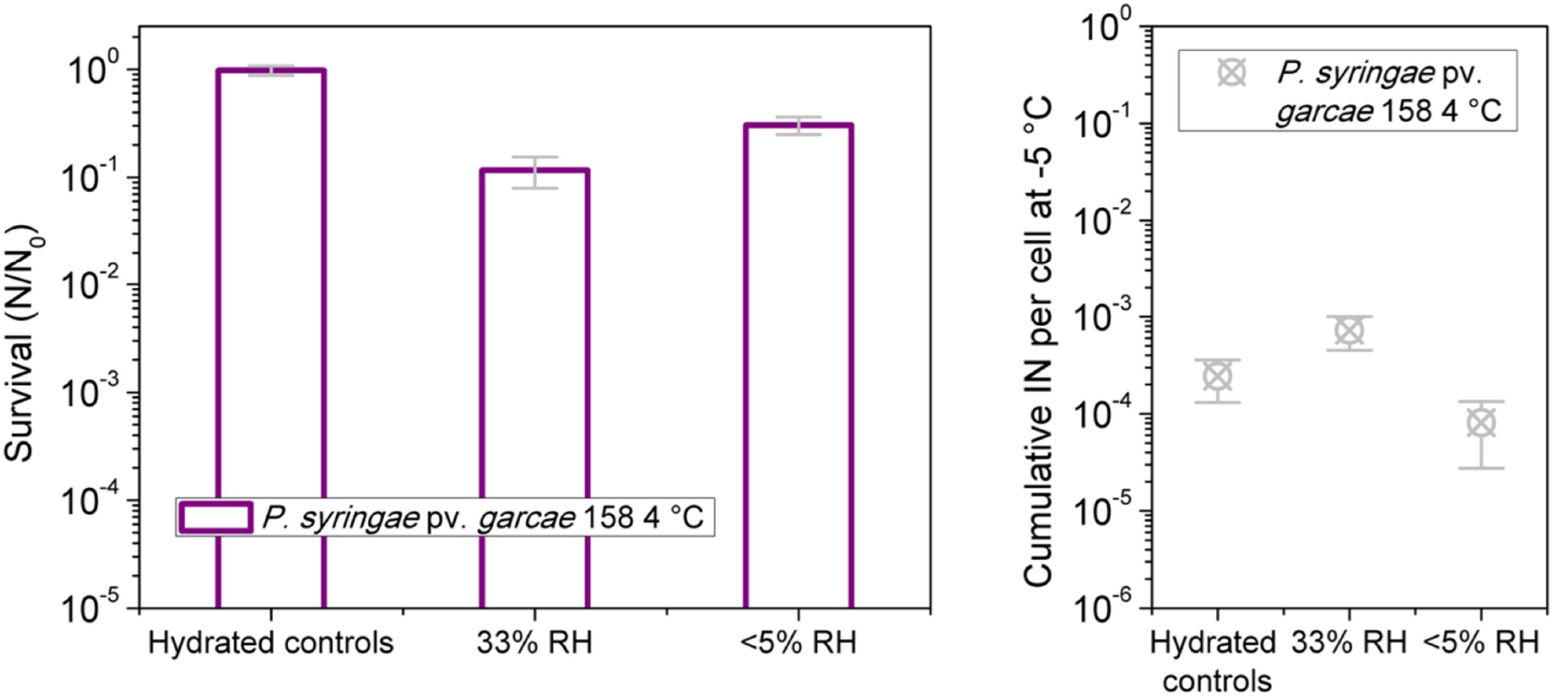
Survival and concentration of cumulative ice nuclei per cell at -5 °C of *P. syringae* pv. *garcae* 158 desiccated at RH 33% and <5% for 6 days at 4 °C in a refrigerator, including its hydrated controls.

